# A rhesus macaque model of Asia lineage Zika virus infection

**DOI:** 10.1101/046334

**Authors:** Dawn M. Dudley, Matthew T. Aliota, Emma Mohr, Andrea M. Weiler, Gabrielle Lehrer-Brey, Kim L. Weisgrau, Mariel S. Mohns, Meghan E. Breitbach, Mustafa N. Rasheed, Christina M. Newman, Dane D. Gellerup, Louise H. Moncla, Jennifer Post, Nancy Schultz-Darken, Michele L. Schotkzo, Jennifer M. Hayes, Josh A. Eudailey, M. Anthony Moody, Sallie R. Permar, Shelby L. O’Connor, Eva G. Rakasz, Heather A. Simmons, Saverio Capuano, Thaddeus G. Golos, Jorge E. Osorio, Thomas C. Friedrich, David H. O’Connor

## Abstract

Infection with Asian lineage Zika virus has been associated with Guillain-Barré syndrome and fetal abnormalities ^1–4^, but the mechanisms and risk factors for these outcomes remain unknown. Here we show that rhesus macaques are susceptible to infection by an Asian-lineage virus closely related to strains currently circulating in the Americas. Following subcutaneous inoculation, Zika virus RNA was detected in plasma one day post-infection (dpi) in all animals (N = 8, including 2 animals infected during the first trimester of pregnancy). Plasma viral loads peaked above 1 × 10^5^ viral RNA copies/mL in seven of eight animals. Viral RNA was also present in saliva, urine, and cerebrospinal fluid (CSF), consistent with case reports from infected humans. Viral RNA was cleared from plasma and urine by 21 dpi in non-pregnant animals. In contrast, both pregnant animals remained viremic longer, up to 57 days. In all animals, infection was associated with transient increases in proliferating natural killer cells, CD8+ T cells, CD4+ T cells, and plasmablasts. Neutralizing antibodies were detected in all animals by 21 dpi. Rechallenge of three non-pregnant animals with the Asian-lineage Zika virus 10 weeks after the initial challenge resulted in no detectable virus replication, suggesting that primary Zika virus infection elicits protective immunity against homologous virus strains. These data establish that Asian-lineage Zika virus infection of rhesus macaques provides a relevant animal model for studying pathogenesis in pregnant and non-pregnant individuals and evaluating potential interventions against human infection, including during pregnancy.

---

Zika virus (ZIKV) is a mosquito-borne flavivirus first identified in 1947^5^. Little was known about ZIKV when fetal abnormalities and Guillain-Barré syndrome were reported coincident with epidemic spread of Asian-lineage ZIKV in South America^6,7,1^. Animal models are essential for quickly understanding ZIKV transmission and pathogenesis, as well as for evaluating candidate vaccines and therapeutics. ZIKV infects immunocompromised mice^8^ but such mice do not mimic key attributes of human infection and fetal development.

In contrast, immunocompetent macaque monkeys are widely used in both infectious disease and obstetric research. To determine whether a phyisiologically relevant dose and route of Asian lineage ZIKV infects immunocompetent pregnant and non-pregnant macaques, we inoculated eight Indian-origin rhesus macaques *(Macaca mulatta)* subcutaneously with ZIKV derived from a French Polynesian virus isolate (Zika virus/H.sapiens-tc/FRA/2013/FrenchPolynesia-01_v1c1). The eight animals were divided into three cohorts as shown in Extended Data Fig. 1.

To define the minimal dosage necessary to establish infection, two macaques per group were infected with 1 x 10^6^, 1 x 10^5^, or 1 x 10^4^ PFU ZIKV (cohorts 1 and 2) (Fig. 1a). This dose range of inocula is based on previous work in related flaviviruses such as West Nile virus (WNV) and DENV, where it was estimated that mosquitoes delivered 1 x 10^4^- 1x 10^6^ PFU of virus^9,10^. This is also the range found in mosquito saliva in a recent publication specifically for Brazilian Zika virus ^11^. None of the animals developed significant clinical disease (see Extended Data Fig. 2 and SI Data)

**Figure 1.**
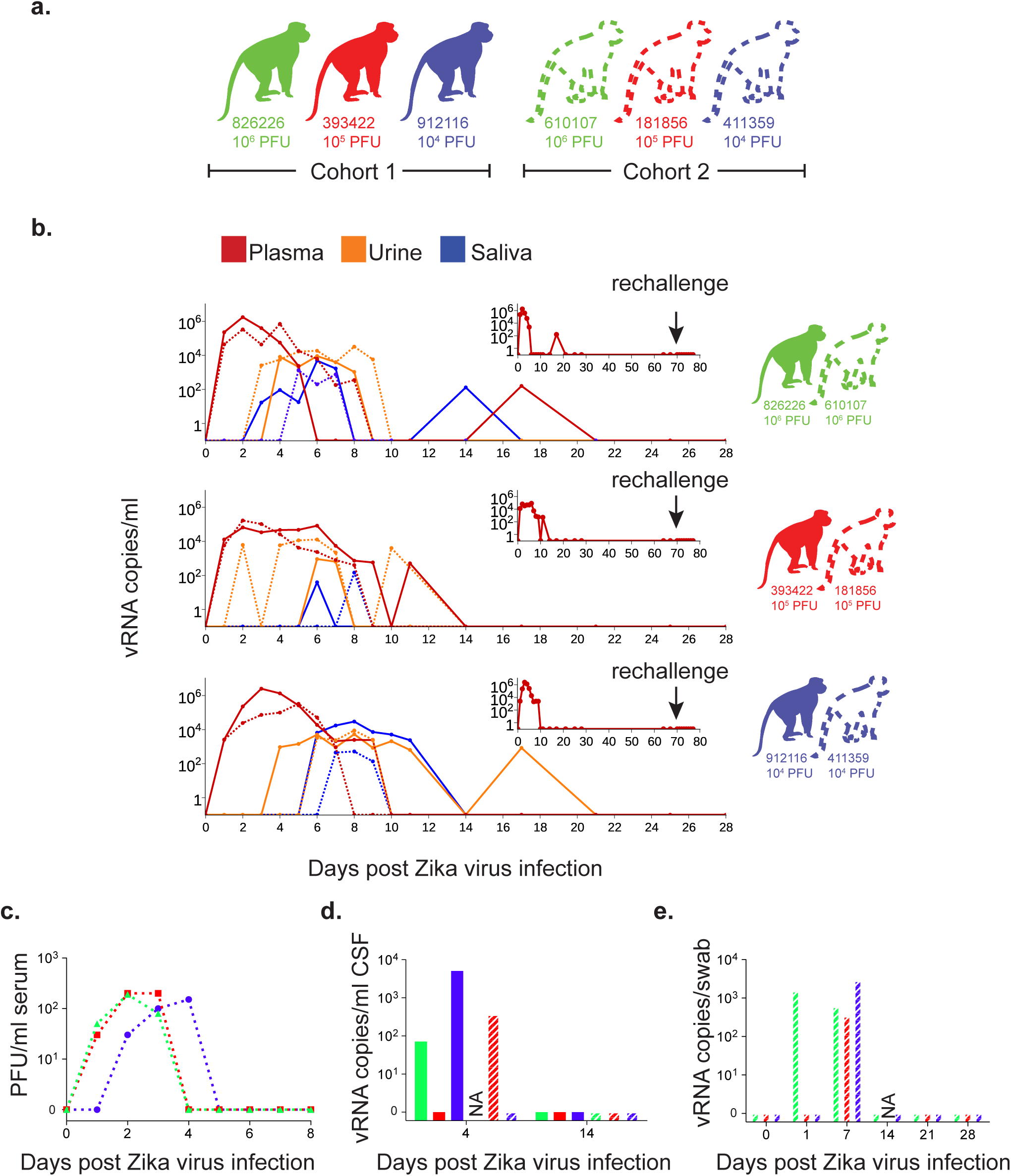
Animal cohort definitions and ZIKV viral load from rhesus macaque fluids. **a.** Animals included in this study and the ZIKV doses used to infect them. Solid lines and bars throughout the figure represent cohort 1 animals by color while stripped bars and dotted lines represent cohort 2 animals by color. **b.** Viral RNA loads measured in plasma, urine, and saliva for the two animals challenged with each dose of virus through 28 dpi. Cohort 1 animals are represented by a solid line while cohort 2 animals are represented by a dotted line for each fluid. Inset: vRNA loads from cohort 1 animals measured before and after rechallenge with homotypic Zika virus as indicated by an arrow. **c.** Number of plaque forming units per ml of serum for cohort 2 animals. **d.** Viral RNA load per ml of CSF collected on 4 and 14 dpi. e. Viral RNA load per vaginal swab collected on 0, 7, 14, 21 and 28 dpi. NA: sample not available.

Blood was sampled daily for 10-11 days post-infection (dpi) and every 3-7 days thereafter. Viral RNA (vRNA) was quantified by qRT-PCR from plasma^12^ and was detected in all six animals at 1 dpi (Fig. 1b). Peak plasma viremia occurred between 2 and 6 dpi, and ranged from 8.2 x 10^4^ to 2.5 x 10^6^ vRNA copies/mL. These results resemble findings in humans in Colombia where the mean serum viral load was 2.6 x10^5^ copies/ml ± 10 copies/ml in acutely infected individuals (n=10, range=537 - 6.9 x 10^5^ copies/ml) (manuscript submitted^13^). Infectious titers, measured from serum in cohort 2 animals, were 500-1000-fold less than copies of vRNA detected from plasma at the same time points (Fig. 1c). Copies of vRNA detected in the serum and plasma were very similar as shown in Extended Data Fig. 3. The estimated doubling time for plasma viremia averaged 7.7 hours (range = 4.8-10.2 hours) and was independent of the infecting dose and sex of the macaque. By 10 dpi, plasma viral loads were undetectable (<100 vRNA copies/mL) in all six animals, although intermittent low-level detection (<550 vRNA copies/mL) continued sporadically through 17 dpi. Thereafter viral RNA remained undetectable in all fluids throughout follow-up (longest follow-up 70 dpi; Fig. 1b insets).

We also measured ZIKV vRNA by qRT-PCR in other body fluids including urine, saliva, CSF and vaginal fluid. Viruria was detected starting at 2-5 dpi and as late as 17 dpi, in urine passively collected from cage pans (Fig. 1b). Despite possible degradation of virus between the time of urination and sample collection and processing, 1 x 10^3^-1 x 10^4^ vRNA copies/mL urine were detected at multiple timepoints. Virus was also detected in oral swabs collected from all six animals, peaking at over 1 x 10^3^ vRNA copies per sample in 3 of 6 animals (Fig. 1b). Notably, as with urine, the kinetics of virus detection in saliva occurred after peak plasma viremia. Cisterna magna punctures were performed at 4 and 14 dpi to quantify viral RNA in CSF; vRNA was detectable at 4 dpi in 3 out of 5 animals from which CSF could be obtained (Fig. 1d). Vaginal swabs collected from the 3 female animals in cohort 2 had detectable vRNA starting at 1 and/or 7 dpi, but were undetectable at 14, 21, and 28 dpi (Fig. 1e).

We next characterized the immune response to infection by staining peripheral blood mononuclear cells (PBMC) for multiple lineage and activation markers. Proliferating (Ki-67+) NK cells, CD8+ T cells, and CD4+ T cells expanded significantly above baseline levels by 6 dpi (Fig. 2a,b). NK and CD8+ T cell expansion increased as plasma vRNA loads decreased starting at 6 dpi. We also enumerated circulating plasmablasts, defined as CD3-/20-/14-/16-/11c-/123- and CD80+/HLA-DR+ cells, on 0, 3, 7, 11 and 14 dpi (Fig. 2c)^14^. The peak plasmablast expansion occurred between 7 and 10 dpi in 5/6 animals. Serum neutralizing antibody responses were also measured by plaque reduction neutralization tests (PRNT_90_). All animals exhibited high neutralizing antibody (nAb) titers as early as 14 dpi (Fig. 2d), the earliest time point tested. Cohort 1 animals were tested at 64 dpi and cohort 2 animals were tested at 14 and 28 dpi. Together these data suggest that peak activation of the adaptive immune response and antibody production occurs 5-7 dpi and may both be important to control viral replication as evidenced by reducing vRNA loads in the plasma at these time points.

**Figure 2.**
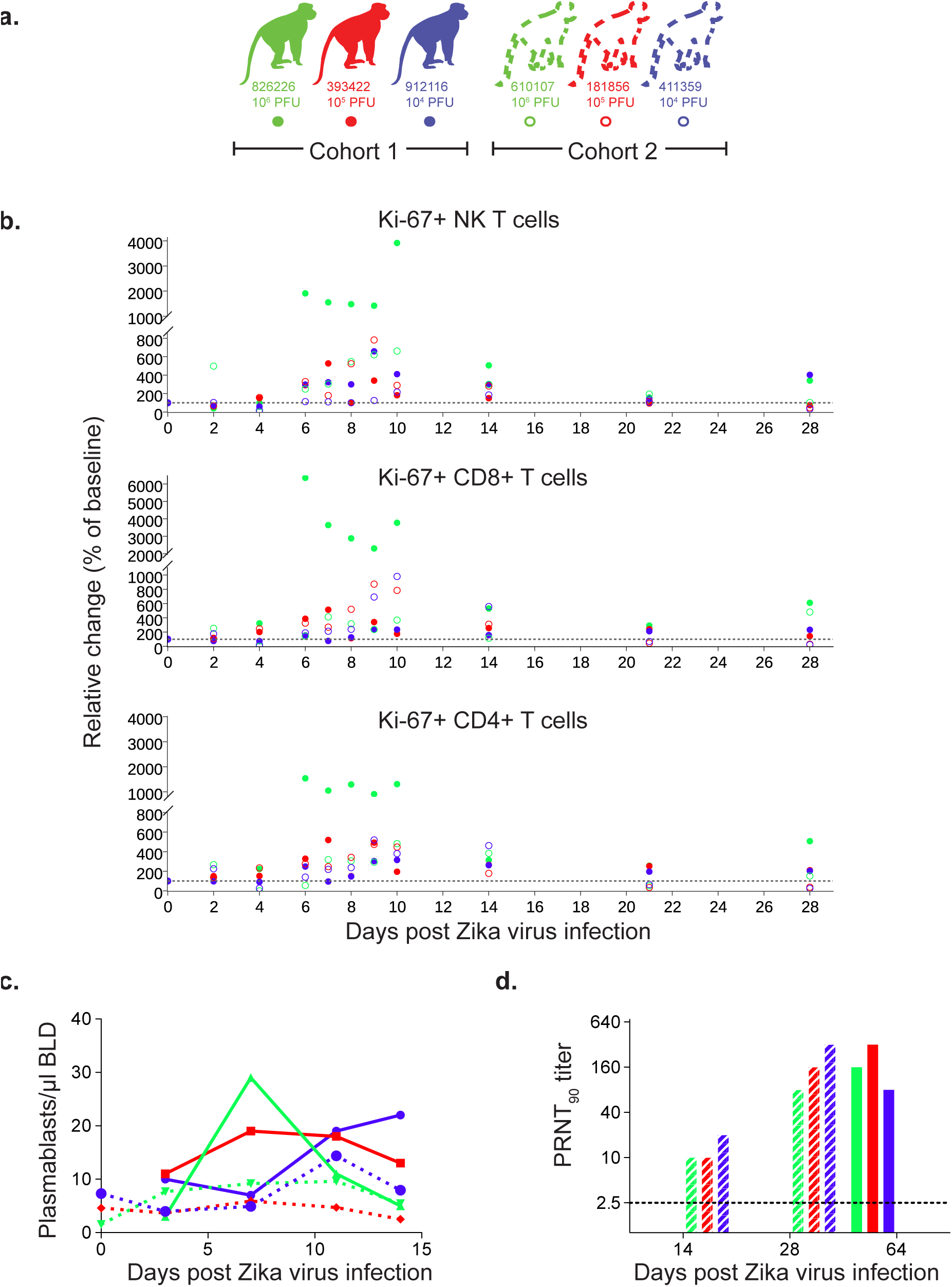
Immune cell expansion and neutralizing antibody titers following ZIKV infection. a. Solid dots, lines and bars with corresponding color represent cohort 1 animals and open circles, dotted lines or stripped bars represent cohort 2 animals throughout the figure. **b.** Expansion of Ki-67+ (activated) NK cells, CD8+ T cells and CD4+ T cells were measured daily for 10 days and then on days 14, 21 and 28 post-infection. Absolute numbers of activated cells/μl of blood are presented relative to the baseline value set to 100%. **c.** Total number of plasmablast cells found in PBMCs collected at 0 (cohort 2 only), 3, 7, 11 and 14 dpi for each animal. **d.** PRNT_90_ titers for cohort 1 and cohort 2.

To determine whether activation of T cells correlated with the appearance of Zika virus-specific responses, we performed interferon-gamma ELISPOT on PBMCs collected at 4, 10 and 14 dpi for cohort 2 animals. Cells were stimulated with pools of 15mer peptides collectively representing the amino acid sequence of the Asian-lineage NS5 protein (GenBank: KU321639). We detected specific IFN-gamma secretion in response to 12 of 16 peptide pools in at least one animal (Extended Data Fig. 4a). Overall, this data supports that there are ZIKV-specific T cell responses in all animals tested.

To determine whether the immune responses that we detected following primary challenge were protective against homotypic rechallenge, we rechallenged the three animals in cohort 1 ten weeks after primary infection with 1 x 10^4^ PFU of a homologous virus (Fig. 1b inset and Extended Data Fig. 1). Plasma, urine and saliva vRNA loads remain negative to at least 9 dpi (as of 5/4/2016), indicating complete protection against ZIKV re-infection.

We also challenged two time-mated rhesus macaques at approximately gestation day 31 and 38 (mid-first trimester) with 1 x 10^4^ PFU of ZIKV (cohort 3; see Extended Data Fig. 1 and Fig. 3a). Both animals were viremic by 1 dpi and exhibited peak plasma viral loads of > 4 x 10^5^ vRNA copies/ml by 3 or 6 dpi (Fig. 3b). Infectious virus was also quantitated by plaque assay from the serum of 660875 (Extended Data Fig. 5). In contrast to their non-pregnant counterparts, both animals maintained persistent plasma viremia (vRNA copies/ml) to 57+ and 29 dpi (Fig. 3b). This is similar to a case described by Driggers et. al., where a pregnant mother had persistent ZIKV vRNA detected from 35 to 70 dpi that did not resolve until termination of pregnancy^7^. The fetus was found to have 2x10^8^ copies/ml of virus in brain tissue and it is speculated that the fetus may have been the source of the prolonged maternal plasma viremia. We will continue to monitor these pregnant animals for vRNA in the blood and amniotic fluid and will determine the infection status of the fetus upon termination of the pregnancy either at full term or earlier if necessary for the health and safety of the mother. Amniocentesis using ultrasound guidance was performed at 43 dpi for 827577 and 36 dpi for 660875 and both were negative for ZIKV RNA.

**Figure 3.**
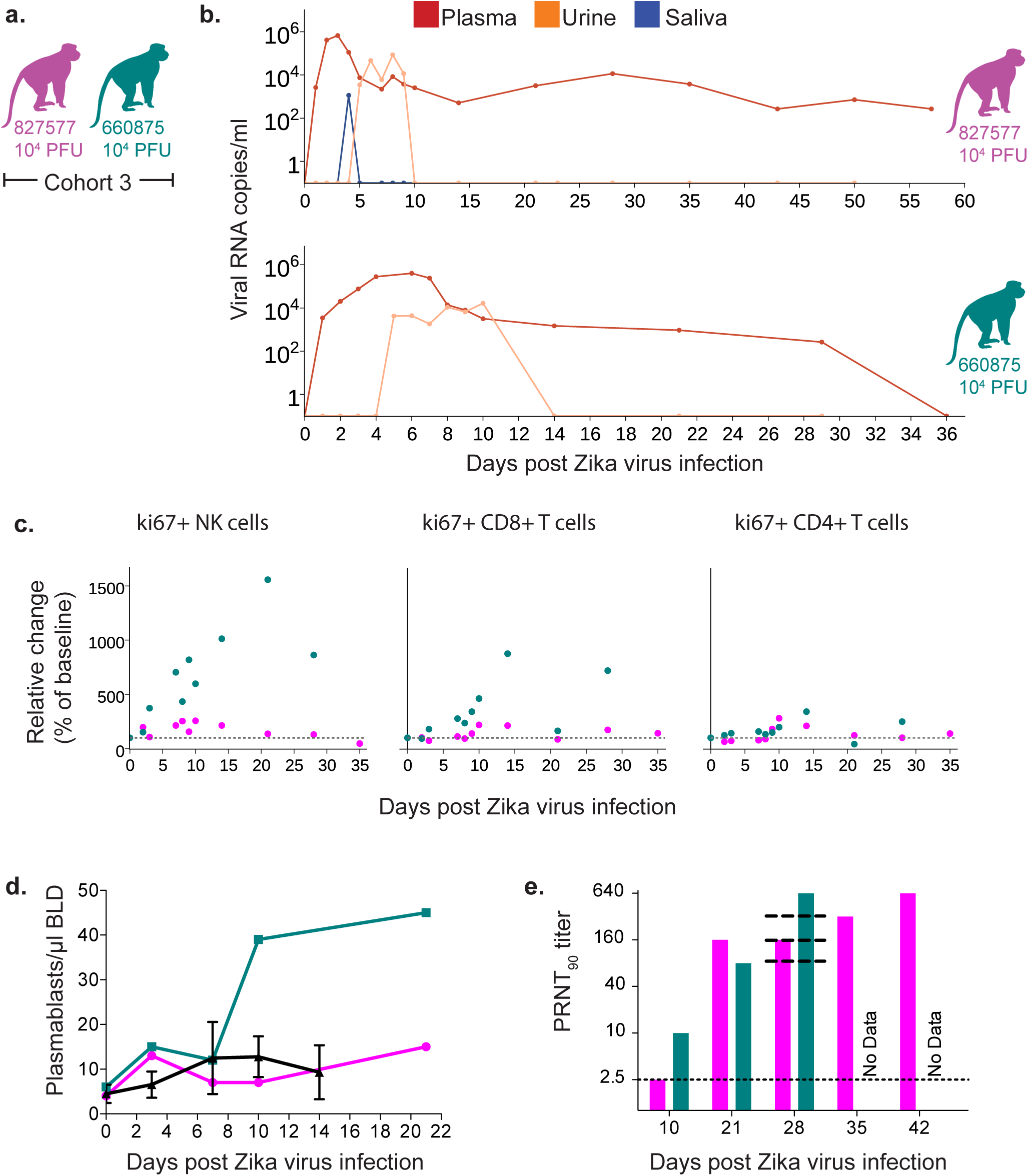
Characterization of Zika virus infection in animals infected during the first trimester of pregnancy. **a.** Schematic of animals presented in this figure as cohort 3. Dots, lines and bars representing each animal match in color throughout the figure. **b.** vRNA copies of the plasma, urine, and saliva from each pregnant animal. Oral swabs could not be obtained from 660875. **c.** Absolute numbers of Ki-67+ NK, CD8+ T cell and CD4+ T cell populations presented as a percentage relative to baseline (x100) over time in each animal. **d.** Plasmablast expansion over time from each pregnant animal. The average plasmablast expansion of cohort 1 and cohort 2 animals infected with the 10^4^ PFU is presented by the black line. Error bars represent standard deviation. **e.** PRNT_90_ titers over time for each animal. Lines representing the titers from cohort 2 animals are overlayed at 28 dpi for reference (top to bottom: 610107, 181856, and 411359).

Both animals generated similar activation of NK, CD8+ T cell and CD4+ T cell responses above baseline as non-pregnant animals (Fig. 3c). Expansion of plasmablast cells was also observed by 10-21 dpi with one animal expanding more than, and one animal expanding less than, the average non-pregnant animal (Fig. 3d). Neutralizing antibodies were detected by 21 dpi for 827577 and 10 dpi for 660875 and were similar to the cohort 2 non-pregnant animals at 28 dpi (Fig. 3e). One pregnant animal exhibited an increase in CK above RI and above levels expected with repeated ketamine sedation and blood collection. The same pregnant animal also developed persistent regenerative anemia characterized by circulating nucleated erythrocytes. Updates to the cohort 3 experiments are available in real-time at goo.gl/rmNCqf.

Altogether, our study shows the persistence of ZIKV RNA in the plasma of rhesus macaques for approximately 10 days, similar to other vector-borne flaviviruses that cause acute, typically self-limiting infections in humans. This work also shows that natural ZIKV infection elicits a robust immune response including ZIKV-specific T cell response and neutralizing antibody responses that confers protection against reinfection. However, the prolonged detection of vRNA in urine and saliva after apparent clearance from the blood, detection of virus in the CSF, and occasional plasma “blips” after initial clearance, suggest that ZIKV may persist longer, at low levels, in certain tissues. Future work in rhesus macaques will seek to determine if and where these reservoirs may exist and whether they seed virus into fluids that might allow for human-to-human transmission.

Our study establishes immunocompetent rhesus macaques infected with physiologically relevant ZIKV as a relevant translational model for infection and pathogenesis. The large immunological toolset available for rhesus macaques will enable investigations of immunity and potential vaccines. Pregnancy, the maternal-fetal interface, and fetal development have been described in detail in rhesus macaques, so this model will also enable assessments of the impact of maternal ZIKV infection on the developing fetus. We have established persistent viremia in pregnant macaques despite activation of NK cells and T cells as well as development of a neutralizing antibody response. We continue to follow these pregnant animals and will establish whether fetal infection and/or abnormalities have occurred through serial ultrasound assessments of the fetus and placenta as well as tissue analysis at pregnancy termination.

## Methods

### Study design

This was a proof-of-concept study designed to establish the infectivity and viral dynamics of Asian lineage ZIKV. Because nothing is known about ZIKV dosing in macaques, one male and one female rhesus macaque of Indian ancestry were each challenged with the following ZIKV doses: 1x10^6^, 1x10^5^, and 1x10^4^ PFU ZIKV. Two pregnant macaques at 31 and 38 days of gestation were infected with 1x10^4^ PFU ZIKV. We selected 2 animals per inoculum dose and 2 pregnant animals as a minimum number of animals for this pilot study to provide proof-of-concept and design larger studies necessary to place statistical significance on the findings. All macaques utilized in the study were free of Macacine herpesvirus 1, Simian Retrovirus Type D, Simian T-lymphotropic virus Type 1, and Simian Immunodeficiency Virus. Primary data from the study is available at goo.gl/rmNCqf.

### Care and use of macaques at the Wisconsin National Primate Research Center

All macaque monkeys used in this study were cared for by the staff at the Wisconsin National Primate Research Center (WNPRC) in accordance with the regulations and guidelines outlined in the Animal Welfare Act and the Guide for the Care and Use of Laboratory Animals and the recommendations of the Weatherall report. This study was approved by the University of Wisconsin-Madison Graduate School Institutional Animal Care and Use Committee (Animal Care and Use Protocol Number G005401). For all procedures (i.e., physical examinations, virus inoculations, ultrasound examinations, blood and swab collection), animals were anesthetized with an intramuscular dose of ketamine (10 mL/kg). Blood samples were obtained using a vacutainer system or needle and syringe from the femoral or saphenous vein.

### Inoculations

ZIKV strain H/PF/2013 (GenBank: KJ776791), originally isolated from a 51-year-old female in France returning from French Polynesia with a single round of amplification on Vero cells, was obtained from Xavier de Lamballerie (European Virus Archive, Marseille France). We deep sequenced the challenge stock to verify the expected origin. The ZIKV challenge stock consensus sequence matched the Genbank sequence (KJ776791) of the parental virus, but there were 8 sites where between 5-40% of sequences contained variants that appear to be authentic (6/8 were non-synonymous changes) (Extended Data Fig. 6). Sequences have been deposited in the Sequence Read Archive (SRA) under accession number SRP072852.

Virus stocks were prepared by inoculation onto a confluent monolayer of C6/36 mosquito cells. These cell lines were obtained from ATCC, were not further authenticated and were not specifically tested for mycoplasma. A single harvest of virus with a titer of 1.26 x 10^6^ PFU/mL (equivalent to 1.43 x 10^9^ vRNA copies/mL) was used for all 8 challenges. The stock was thawed, diluted in PBS to the appropriate concentration for each challenge, and loaded into a 1 mL syringe that was kept on ice until challenge. Animals were anesthetized as described above, and 1 mL of inocula was administered subcutaneously over the cranial dorsum. Post-inoculation, animals were closely monitored by veterinary and animal care staff for adverse reactions and signs of disease.

### Viral RNA isolation from plasma

Fresh plasma and PBMC were isolated from EDTA-treated whole blood by Ficoll density centrifugation at 1860 rcf for 30min. The plasma layer was collected and centrifuged for an additional 8 min at 670 rcf to remove residual cells. RNA was extracted from 300 μl of plasma using the Viral Total Nucleic Acid Purification Kit (Promega, Madison, WI) on a Maxwell 16 MDx instrument. The RNA was then quantified by quantitative RT-PCR.

### Viral RNA isolation from urine

Urine was collected from a pan beneath the animal’s cage. Urine was centrifuged for 5 min at 500 rcf to remove cells and other debris. RNA was isolated from 300 μl urine using the Viral Total Nucleic Acid Purification Kit (Promega, Madison, WI) on a Maxwell 16 MDx instrument.

### Viral RNA isolation from oral swabs

Oral swab samples were collected from infected animals while anesthetized by gently running a sterile swab under the animal’s tongue. Swabs were placed immediately into either RNAlater or viral transport medium (tissue culture medium 199 supplemented with 0.5% FBS and 1% antibiotic/antimycotic) for 60-90 minutes. Samples were vortexed vigorously, then centrifuged for 5 min at 500 rcf before removing the swabs. Samples were stored at either ‐20°C (RNAlater samples) or ‐80°C (viral transport medium) until processing. Prior to extraction, virus was pelleted by centrifugation for 1 hour at 4°C at 14000 rpm. Supernatant was removed, leaving the virus in 200 μl media. Viral RNA was extracted from these samples using the Qiamp MinElute Virus Spin kit (Qiagen, Germantown, Maryland) with all optional washes. Viral load data from oral swabs are expressed as vRNA copies/mL eluate.

### Quantitative reverse transcription PCR (qRT-PCR)

Viral RNA isolated from plasma, urine, or oral swabs was quantified by qRT-PCR using the primers and probe designed by Lanciotti et al. ^12^.The RT-PCR was performed using the SuperScript III Platinum one-step quantitative RT-PCR system (Invitrogen, Carlsbad, CA) on the LightCycler 480 instrument (Roche Diagnostics, Indianapolis, IN). Primers and probe were used at final concentrations of 600 nm and 100 nm respectively, along with 150 ng random primers (Promega, Madison, WI). Cycling conditions were as follows: 37°C for 15 min, 50°C for 30 min and 95°C for 2 min, followed by 50 cycles of 95°C for 15 sec and 60°C for 1 min. Virus concentration was determined by interpolation onto an internal standard curve composed of seven 10-fold serial dilutions of a synthetic ZIKV RNA fragment based on the Asian-lineage (ZIKV strain H/PF/2013).

### Viral quantification by plaque assay

Titrations for replication competent virus quantification of the challenge stock as well as from serum collected at multiple time points from cohort 2 animals were completed by plaque assay on Vero cell cultures. Vero cells were obtained from ATCC, were not further authenticated and were not specifically tested for mycoplasma. Duplicate wells were infected with 0.1 mL aliquots from serial 10-fold dilutions in growth media and virus was adsorbed for one hour. Following incubation, the inoculum was removed, and monolayers were overlaid with 3 ml containing a 1:1 mixture of 1.2% oxoid agar and 2X DMEM (Gibco, Carlsbad, CA) with 10% (vol/vol) FBS and 2% (vol/vol) penicillin/streptomycin. Cells were incubated at 37°C in 5% CO_2_ for four days for plaque development. Cell monolayers then were stained with 3 mL of overlay containing a 1:1 mixture of 1.2% oxoid agar and 2X DMEM with 2% (vol/vol) FBS, 2% (vol/vol) penicillin/streptomycin, and 0.33% neutral red (Gibco). Cells were incubated overnight at 37°C and plaques were counted. Titers of virus detected from the serum of cohort 2 animals were compared to plasma and serum viral load assays. For both the challenge stock and the virus isolated from macaque serum, the level of infectious virus detected by plaque assay was ~500-1000-fold less than the number of viral RNA particles detected by qRT-PCR in either the plasma or serum. This was true throughout the duration of viremia where plaque assay titers were detectable.

### Plaque reduction neutralization test (PRNT_90_)

Macaque serum samples were screened for ZIKV neutralizing antibody utilizing a plaque reduction neutralization test (PRNT). Endpoint titrations of reactive sera, utilizing a 90% cutoff (PRNT90) were performed as described^15^ against ZIKV strain H/PF/2013.

### Immunophenotyping

The amount of activated/proliferating NK cells were quantified using a modified version of our protocol detailed step-by step in OMIP-28^16^. Briefly 0.1 mL of EDTA-anticoagulated whole blood samples were incubated for 15 min at room temperature in the presence of a mastermix of antibodies against CD45 (clone D058-1283, Brilliant Violet 786 conjugate), CD3 (clone SP34-2 Alexa Fluor 700 conjugate), CD8 (clone SK2, Brilliant Violet 510), NKG2A/C (clone Z199, PE-Cy7 conjugate), CD16 (clone 3G8, Pacific Blue conjugate), CD69 (clone TP1.55.3, ECD conjugate), HLA-DR (clone 1D11, Brilliant Violet 650 conjugate), CD4 (clone SK3, Brilliant Violet 711 conjugate), CCR7 (clone 150503, Fluorescein conjugate), CD28 (clone CD28.2, PE conjugate), and CD95 (clone DX2, PE-Cy5 conjugate) antigens. All antibodies were obtained from BD BioSciences, except the NKG2A/C-specific antibody, which was purchased from Beckman Coulter, and the CCR7 antibody that was purchased from R&D Systems. Red blood cells were lysed using BD Pharm Lyse, after which they were washed twice in media and fixed with 0.125 ml of 2% paraformaldehyde for 15 min. After an additional wash the cells were permeabilized using Life Technology’s Bulk Permeabilization Reagent. The cells were stained for 15 min. with Ki-67 (clone B56, Alexa Fluor 647 conjugate) while the permeabilizer was present. The cells were then washed twice in media and resuspended in 0.125 ml of 2% paraformaldehyde until they were run on a BD LSRII Flow Cytometer. Flow data were analyzed using Flowjo version 9.8.2.

### Interferon-gamma ELISPOT assay

Peripheral blood mononuclear cells (PBMCs) were isolated from EDTA-treated whole blood by using Ficoll-Paque Plus (GE Health Sciences) density centrifugation. Enzyme-linked immunosorbent spot (ELISPOT) assays were conducted according to the manufacturer’s protocol. Briefly, 1 x 10^5^ cells in 100ul of R10 medium were added to pre-coated monkey gamma interferon (IFN_γ_) ELISpot-PLUS plates (Mabtech Inc., Mariemont, OH) with peptide at a final concentration of 1uM. Full proteome peptides derived from the ZIKV NS5 sequence (GenBank: KU321639.1) used in this study were synthesized by GenScript (Piscataway, NJ). Pools were created using 10 overlapping 15mer peptides, each at a working concentration of 1mM. Concanavalin A (10uM) was used as a positive control. Assays of all samples were repeated in duplicate or triplicate. Cells alone in the absence of stimulant were used as a negative control. Wells were imaged by using an AID ELISPOT reader, and spots were counted by using an automated program with parameters including size, intensity, and gradient. The limit of detection was set at 100 spot-forming cells per million PBMCs.

### Plasmablast detection

Peripheral blood mononuclear cells (PBMCs) isolated from three ZIKV-infected rhesus monkeys at 3, 7, 11, and 14 dpi were stained with the following panel of fluorescently labeled antibodies (Abs) specific for the following surface markers: CD20 FITC (L27), CD80 PE(L307.4), CD123 PE-Cy7(7G3), CD3 APC-Cy7 (SP34-2), IgG BV605(G18-145) (all from BD Biosciences, San Jose, CA), CD14 AF700 (M5E2), CD11c BV421 (3.9), CD16 BV570 (3G8), CD27 BV650(O323) (all from BioLegend, San Diego, CA), IgD AF647 (polyclonal)(Southern Biotech, Birmingham, AL), and HLA-DR PE-TxRed (TÜ36) (Invitrogen, Carlsbad, CA). LIVE/DEAD Fixable Aqua Dead Cell Stain Kit (Invitrogen, Carlsbad, CA) was used to discriminate live cells. Briefly, cells were resuspended in 1X PBS/1 %BSA and stained with the full panel of surface Abs for 30 min in the dark at 4°C, washed once with 1X PBS, stained for 30 min with LIVE/DEAD Fixable Aqua Dead Cell Stain Kit in the dark at 4°C, washed once with 1X PBS, washed again with 1X PBS/1%BSA, and resuspended in 2% PFA Solution. Stained PBMCs were acquired on a LSRII Flow Analyzer (BD Biosciences, San Jose, CA) and the data was analyzed using FlowJo software v9.7.6 (TreeStar, Ashland, OR). Plasmablasts were defined similarly to the method previously described^14^ excluding lineage cells (CD14+, CD16+, CD3+, CD20+, CD11c+, CD123+), and selecting CD80+ and HLA-DR+ cells (known to be expressed on rhesus plasmablasts and their human counterpart^17^).

### Estimation of plasma viremia doubling time

The doubling time of plasma viremia was estimated in R version 3.2.3 (The R Foundation for Statistical Computing, http://www.R-project.org). For each animal, the slope of the linear portion of the line (between 1 and 2 dpi for the animals treated with 1x10^6^ and 1x10^5^ PFU and between 1, 2, and 3 dpi for the animal treated with 1x10^4^ PFU) was generated by plotting the log of the plasma viral loads. The linear portion represents the exponential growth phase and has been used to estimate doubling time in other systems^18^. The slopes were then used in the equation: log(2)/slope. Each result was then multiplied by 24 hours to produce a simple estimate of doubling time in hours.

### CBC and blood chemistry panels

CBCs were performed on EDTA-anticoagulated whole blood samples on a Sysmex XS-1000i automated hematology analyzer (Sysmex Corporation, Kobe, Japan). Blood smears were prepared and stained with Wright-Giemsa stain (Wescor Aerospray Hematology Slide Stainer; Wescor Inc, Logan, UT). Manual slide evaluations were performed on samples as appropriate when laboratory-defined criteria were met (including the presence of increased total white blood cell counts, increased monocyte, eosinophil, and basophil percentages, decreased hemoglobin, hematocrit, and platelet values, and unreported automated differential values). Individuals performing manual slide evaluations screened both white blood cells (WBC) and red blood cells (RBC) for cellular maturity, toxic change, and morphologic abnormalities.

Whole blood was collected into serum separator tubes (Becton, Dickinson and Company, Franklin Lakes, NJ) for blood chemistry analysis and processed per manufacturer’s instructions. Blood chemistry panels were performed on the serum using a Cobas 6000 analyzer (Roche Diagnostics, Risch-Rotkreuz, Switzerland). Results from CBC and blood chemistry panels were reported with species, age, and sex-specific reference ranges.

### Zika virus deep sequencing of the challenge stock

A vial of the same ZIKV strain H/PF/2013 virus stock that infected macaques was deep sequenced by preparing libraries of fragmented double-stranded cDNA using methods similar to those previously described^19^. Briefly, the sample was centrifuged at 5000 rcf for 5 min. The supernatant was then filtered through a 0.45-μm filter. The Qiagen QiAmp Minelute viral RNA isolation kit (omitting carrier RNA) was used to isolate vRNA. The eluted RNA was then treated with DNAse I. Double stranded DNA was prepared with the Superscript double stranded cDNA synthesis kit (Invitrogen) and priming with random hexamers. Agencourt Ampure XP beads were used to purify double stranded DNA. The purified DNA was fragmented with the Nextera XT kit (Illumina), tagged with Illumina-compatible primers, and then purified with Agencourt Ampure XP beads. Purified libraries were then sequenced with 2 x 300 bp kits on an Illumina MiSeq. Of note, challenge stock viral loads were 1.43x10^9^ vRNA copies/ml. This results in an input of 7.15 x10^8^ RNA copies into the sequencing reactions. This far exceeds the average depth of coverage of 11,877 (+/-4658) sequences per nucleotide site indicating little resampling effects in our data analysis.

**Supplementary Information** is linked to the online version of the paper at www.nature.com/nature.

## Acknowledgements

We thank the Veterinary, Animal Care, Scientific Protocol Implementation, and the Pathology staff at the Wisconsin National Primate Research Center (WNPRC) for their contribution to this study. We thank the DHHS/PHS/NIH (R01Al116382-01A1 to D.H.O.), (R01Al107157-01A1 to T.G.G.) and (DP2HD075699 to S.R.P.) for funding. We also thank the P51OD011106 awarded to the WNPRC, Madison-Wisconsin. This research was conducted in part at a facility constructed with support from Research Facilities Improvement Program grants RR15459-01 and RR020141-01. The publication’s contents are solely the responsibility of the authors and do not necessarily represent the official views of NCRR or NIH.

## Author contributions

D.H.O., T.C.F., J.E.O., M.T.A., E.M., T.G.G. and D.M.D. designed the experiments. D.H.O., D.M.D., M.T.A., E. M., T.C.F., and L.H.M. drafted the manuscript. M.T.A., and J.E.O. provided and prepared viral stocks and performed plaque assays. A.M.W., G.L-B., and T.C.F. developed and performed viral load assays. K.L.W. and E.G.R. performed immunophenotyping assays. M.S.M., M.E.B., M.N.R., C.M.N., and D.M.D. coordinated and processed macaque samples for distribution. D.D.G., S.L.O., and D.M.D. designed and performed the sequencing experiments. L.H.M. and T.C.F. performed nucleotide diversity calculations. J.P., N.S-D., H.A.S., S.C., and J.M.H. coordinated the macaque infections, sampling, and performed blood chemistries and CBC analysis. M.L.S. coordinated experiments and helped perform ultrasounds on the pregnant macaques. J.A.E., M.A.M., and S.R.P. performed the plasmablast experiments.

## Author information

Sequences have been deposited in the Sequence Read Archive (SRA) under accession number SRP072852. All data from these studies are available at zika.labkey.com. Reprints and permissions information is available at www.nature.com/reprints. The authors declare no competing interests. Correspondence and requests for materials should be addressed to.

**Extended Data Figure 1.**
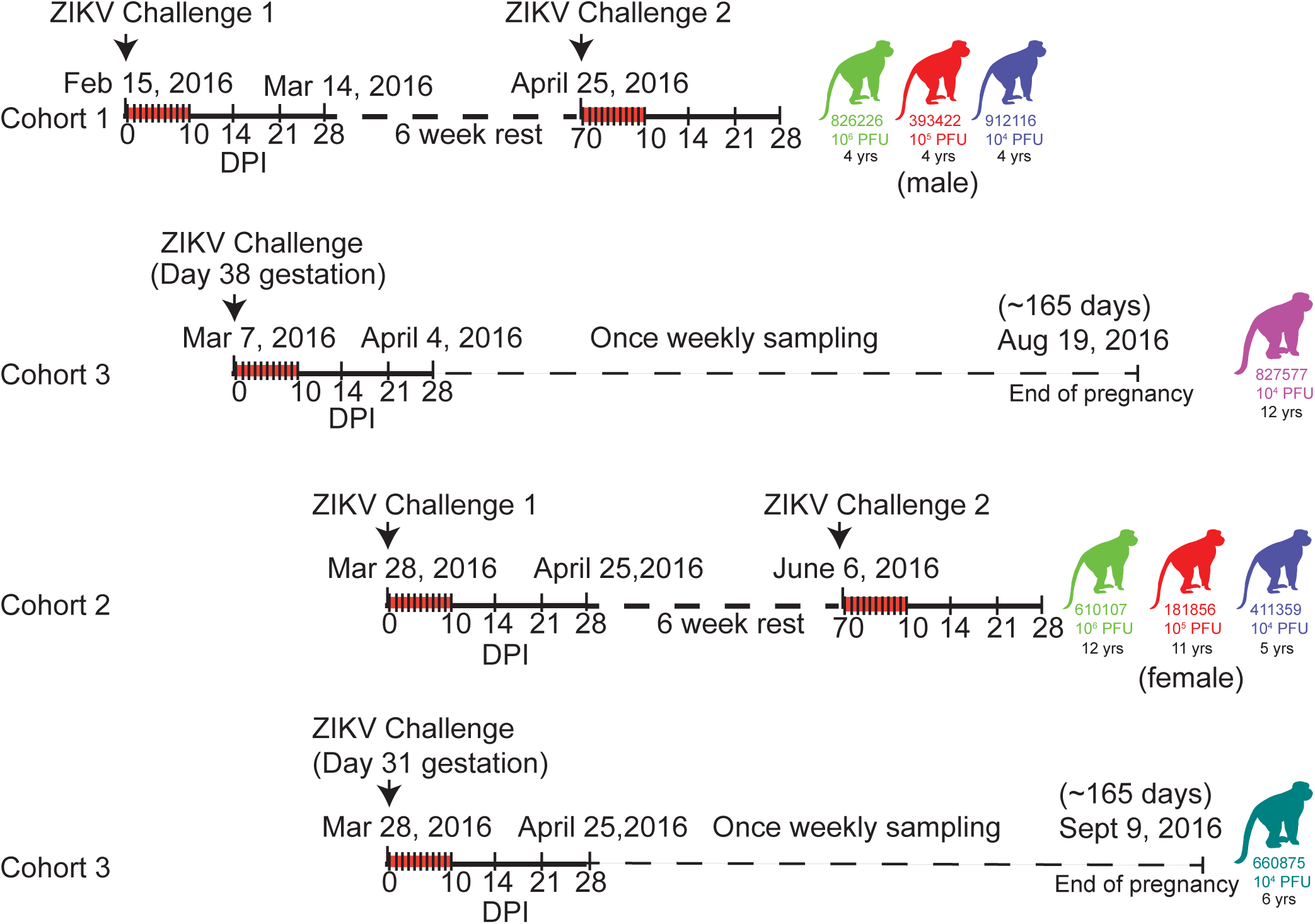
Schematic representation of the timeline of infection and sampling for each animal in the presented studies. Cohort 1 received the first ZIKV challenges and were then rested for 6 weeks before a rechallenge. For all studies, samples were collected daily for 10 days and then on 14, 21, and 28 dpi as indicated by hashes in the timelines. Cohort 3 represents the two pregnant animals that were challenged on 2 different days. Both animals are currently in the once weekly sampling phase until the pregnancies come to term (~165 gestational days). Cohort 2 was a repeat experiment of cohort 1 that allowed for additional experiments and sample collection (e.g., serum plaque infectivity) that were not feasible when we initiated cohort 1 studies. These animals are currently in a 6-week rest period and will be rechallenged on June 6, 2016. Ages of all animals are indicated under each macaque identification number.

**Extended Data Figure 2.**
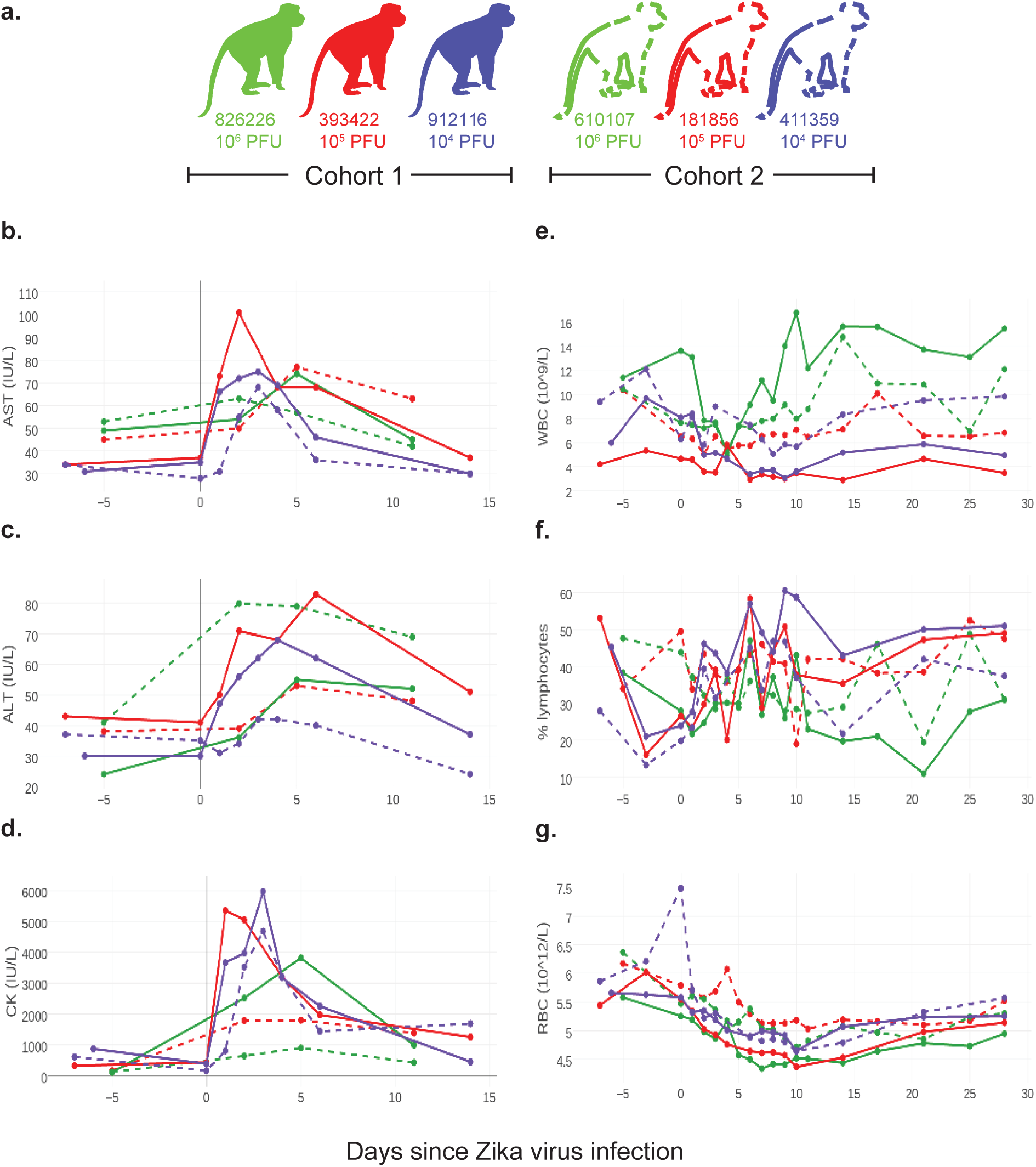
Complete blood counts and serum chemistries for macaques infected with ZIKV. **a.** Animals were infected with different doses of ZIKV. Cohort 1 animals are represented by solid lines and cohort 2 animals are represented by dotted lines. All non-pregnant animals had serum chemistry analysis performed at ‐7, 0, 1, 2, 3, 4, 6, and 14 dpi or at ‐6, 2, 5 and 11 dpi. **b.** AST blood chemistries **c.** ALT serum chemistries. **d.** CK serum chemistries. Complete blood counts were measured prior to infection, daily for 10-11 days after infection and then every 3-7 days until 28 dpi. **e.** white blood cell counts. **f.** % lymphocytes. **g.** red blood cell counts.

**Extended Data Figure 3.**
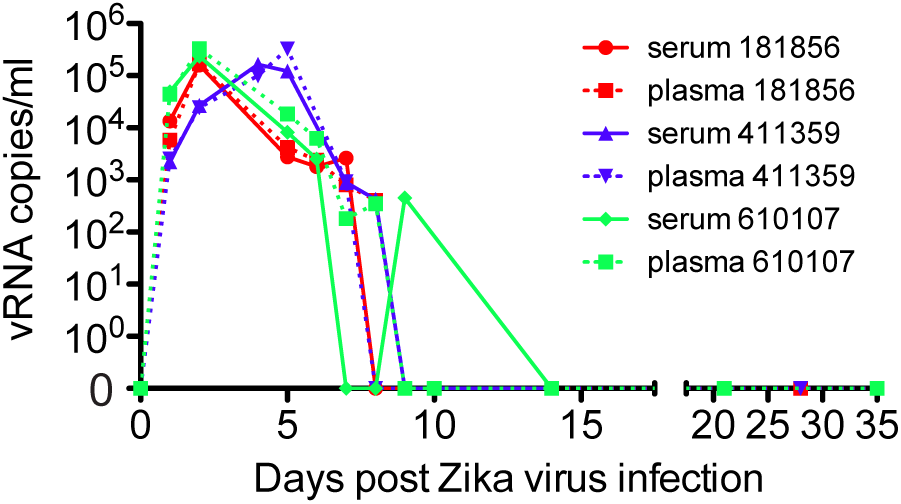
qRT-PCR detection of ZIKV RNA is equally sensitive from serum and plasma. vRNA copies/ml of plasma or serum were quantitated by qRT-PCR over multiple time points from cohort 2 animals. Sufficient baseline samples were not available from serum.

**Extended Data Figure 4.**
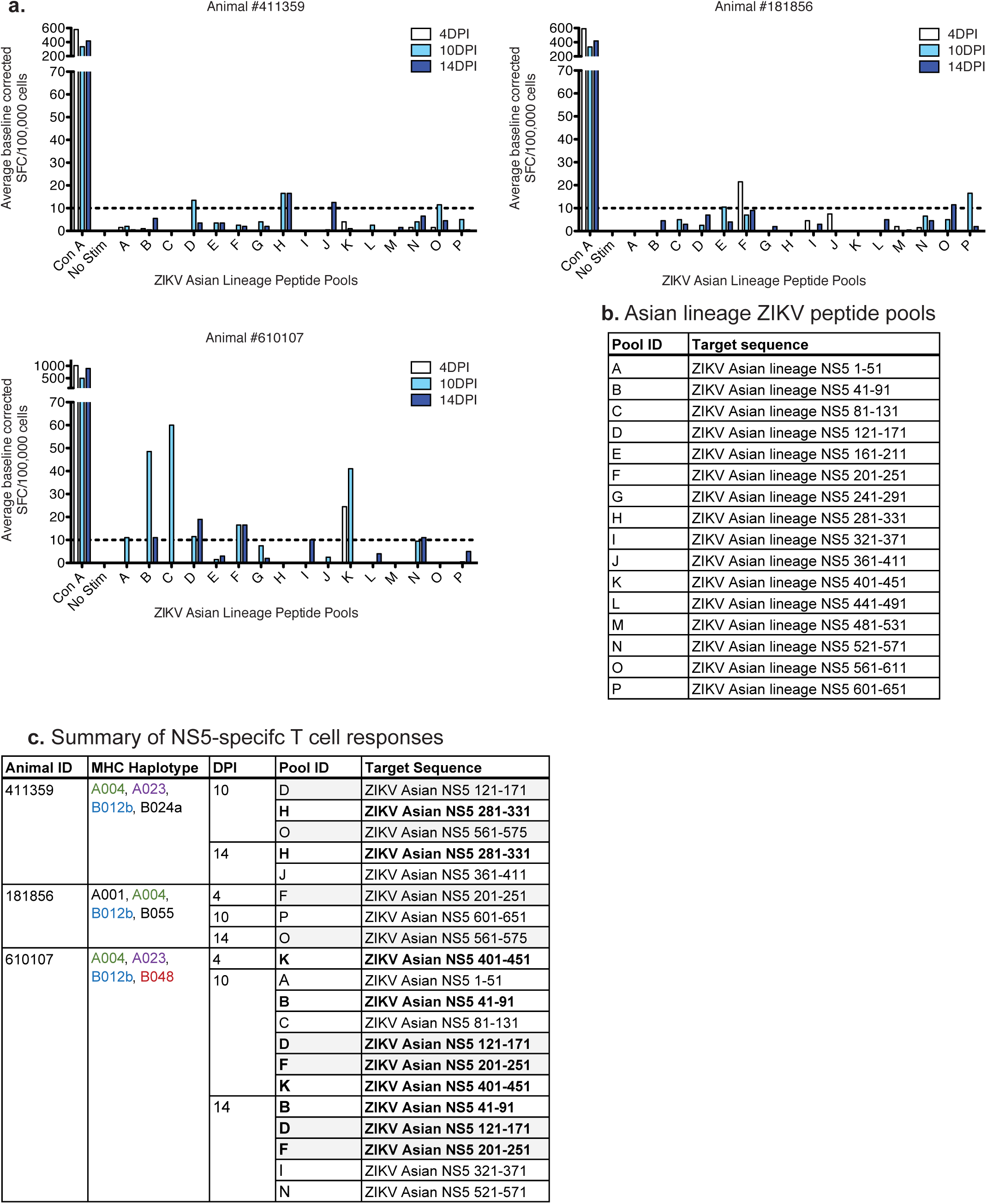
Antigen-specific T cell responses by IFN_γ_-ELISPOT. **a.** Average spot forming cell counts for PBMC collected from each animal at 4, 10 and 14 dpi. Data were baseline corrected by subtracting the average negative control values from each response. A threshold of 10.0 SFC/100,000 cells was set as the minimum value to be considered a positive T cell response, as indicated by the dashed line. **b.** Each pool was comprised of 10 overlapping 15mer peptides offset by 4 amino acids. **c.** Peptide pools eliciting T cell responses at 4, 10 and 14 dpi for each animal. The region of the NS5 protein that is represented by each pool of overlapping 15mers is provided. MHC class I haplotypes of each cohort 2 animal are also presented. All three animals shared the A004 and B012b major histocompatibility complex haplotypes and two animals shared the A023 haplotype. Therefore, it was not surprising that 3 pools were recognized by 2 different animals likely sharing the MHC class I allele that is presenting one of the peptides in those pools. Grayed pools were positive in more than one animal and bolded pools were positive at more than one time point in the same animal.

**Extended Data Figure 5.**
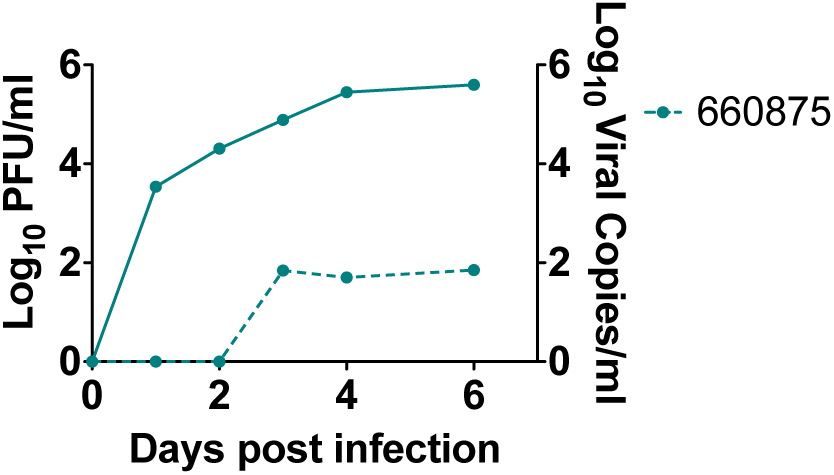
Plaque assay titers in a pregnant animal. Log_10_ PFU/ml serum (dotted line) is plotted relative to vRNA copies/ml plasma (solid line) for 660875.

**Extended Data Figure 6.**
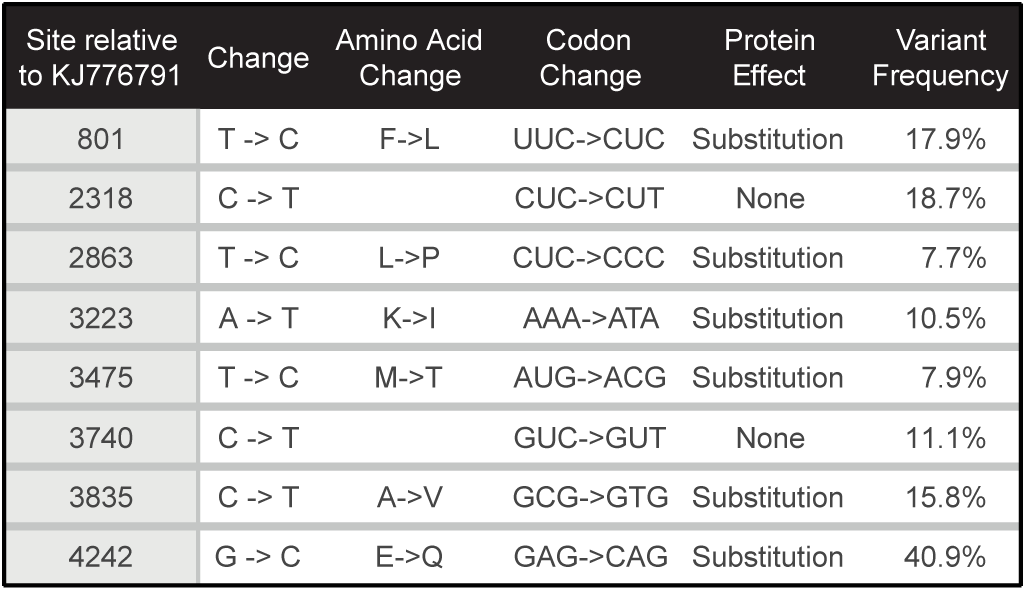
Genetic diversity of the ZIKV challenge stock. The ZIKV challenge stock was deep sequenced from all three animals. Nucleotide sites where at least 5% of sequences obtained from the challenge stock are different from the Genbank sequence are shown.

## SI Guide

### Supplementary Information Data

Morbidity, complete blood counts and serum blood chemistry findings in nonpregnant rhesus macaques exposed to ZIKV. Data is shown in Extended Data Fig. 2.

To detect signs of morbidity, animals were evaluated daily for evidence of disease, injury, or psychological abnormalities (e.g., inappetence, dehydration, diarrhea, depression, inactivity, trauma, self-injurious or stereotypical repetitive behaviors (e.g., pacing) often seen in captive animals). Five of six animals exhibited mild to moderate inappetence, which resulted in mild weight loss in four animals. Two animals (912116 and 393422) also developed a very mild rash around the inoculation site at 1 dpi that persisted for 4-5 days. No other abnormal clinical signs were noted (e.g., increased body temperature, joint pain, lymphadenopathy, lethargy).

Daily complete blood counts (CBCs) were evaluated for all six non-pregnant animals for 10 dpi and then every 3 to 7 days thereafter and serum chemistry analyses were performed intermittently post-infection as per protocol (Extended Data Fig. 2). Reference intervals (RI) developed for the WNPRC colony for species, gender, and age were used to evaluate results. All six animals developed elevated serum creatine kinase (CK), which peaked by 5 dpi (Extended Data Fig. 2d). Increases in serum CK are strongly associated with muscle damage and myositis (skeletal, smooth, and cardiac), but can also be caused by repeated sedation, hemolysis, and endocrine abnormalities^20,21^. Predictable increases in CK have been noted in nonhuman primates undergoing repeated sedation and venipuncture^21^. However, CK levels did not remain elevated during the 10-day period of daily sedation and blood collection, suggesting other causes for the noted increase in values. Future studies are planned to determine if CK increases may be due to viral myositis. ALT values in three of six animals exceeded the maximum WNPRC RI (Extended Data Fig. 2c) and the expected increase associated with repeated ketamine sedation^21^. Although AST values exceeded upper RI values for all six non-pregnant animals after infection, they did not increase above the expected levels previously associated with repeated ketamine sedation described by Lugo-Roman et al. (Extended Data Fig. 2b) ^21^.

All the animals displayed decreased total WBC numbers following infection, but only one animal fell below the RI with a value of 2.88 ths/ul (3.70-15.70). WBC numbers rebounded almost completely to pre-infection levels around 10 dpi in all six animals (Extended Data Fig. 1e). Both animals that received the highest dose of inoculum developed persistent mature neutrophilia around 7-14 dpi that lasted through 28 dpi. Five of 6 non-pregnant animals had mild regenerative anemia characterized by varying degrees of polychromasia and anisocytosis, but whether this was secondary to the viral infection or simply a result of frequent blood collections could not be determined. Platelet values for all six animals remained within RI. Both cellular dyscrasias and elevated transaminases have been described in human ZIKV case reports; myositis has not been reported ^22,23^.

